# Monovalent XBB.1.5 mRNA Vaccine Recalls a More Durable and Coordinated Antibody Response to SARS-CoV-2 Spike than the Bivalent WT/BA.5 mRNA Vaccine

**DOI:** 10.1101/2024.09.12.612662

**Authors:** Susanna E. Barouch, Kate S. Levine, Ross Blanc, Qixin Wang, Xin Tong, Ryan P. McNamara

## Abstract

In the fall of 2023, the monovalent XBB.1.5 mRNA vaccine for COVID-19 became available. However, the comparative magnitude, durability, and functionality of antibody responses induced by the XBB.1.5 vaccine compared with the 2022-2023 bivalent wildtype (WT) + Omicron BA.5 vaccine remains to be fully determined. In this study, we compared antibody profiles generated by these two vaccines in healthcare workers. We show that the monovalent XBB.1.5 vaccine induced higher magnitude binding, neutralizing, and Fc-gamma receptor (FcγR) binding antibodies to the XBB.1.5 spike compared with the bivalent vaccine against the WT and BA.5 spikes, both at both peak immunogenicity and at 6 months post-vaccination. Moreover, antibody interaction architectures and correlations remained more robust at 6 months post-vaccination with the XBB.1.5 vaccine, whereas these correlations were largely lost at 6 months with the bivalent vaccine. Our results suggest that the XBB.1.5 vaccine led to a more durable and functionally coordinated antibody response compared to the bivalent vaccine.

## Introduction

The spike glycoprotein of SARS-CoV-2, the causative agent of coronavirus disease 2019 (COVID-19), is the primary antigen targeted by antibodies generated through infection and/or vaccination. As the virus continues to spread globally, variants of the virus have undergone selective sweeps ^1,2^ with some exhibiting high degrees of escape from antibody-based recognition ^3-9^. This viral evolution has prompted the development and deployment of vaccine boosters with updated spikes to expand antibody recognition, neutralization, and effector function.

Serum neutralizing antibodies have been identified as a correlate of protection against COVID-19 ^10^. These antibodies largely target the receptor binding domain (RBD) of the spike, which binds to the angiotensin-converting enzyme 2 (ACE2) on the surface of respiratory-tract endothelial cells and mediates viral entry ^8,11-15^. Numerous mutations have accumulated in the RBD of virus variants with reduced neutralizing antibody potency. This was observed by the emergence of the Omicron variants ^7,16^. Since the emergence of BA.1, various Omicron sublineages have emerged ^17-19^, including recombinant XBB lineage strains ^20-24^.

The reduced antibody recognition of Omicron variants prompted the deployment of bivalent mRNA vaccines in fall 2022 that encoded both the ancestral spike and the BA.5 spike, which were the predominant forms of the virus at the time of development ^25,26^. Immunologic studies with the bivalent mRNA vaccines demonstrated disproportionate back-boosting to ancestral wildtype (WT) spike and lower responses to the BA.5 spike, potentially as a result of immune imprinting ^25,27-30^. In fall 2023, the monovalent XBB.1.5 mRNA vaccines were deployed.

In this study, we compared antibody responses in recipients of the 2022-2023 bivalent mRNA vaccine (WT + BA.5) with antibody responses in recipients of the 2023-2024 monovalent mRNA vaccine (XBB.1.5) over a period of 6 months. We observed that the monovalent XBB.1.5 booster elicited significantly higher binding and neutralizing antibodies for over 6 months post-immunization compared to the bivalent booster to their respective target antigens. Moreover, the XBB.1.5 booster elicited a sustained and coordinated humoral response, with multiple humoral responses correlated with each other, suggesting that both Fab and Fc regions contributed to neutralizing and non-neutralizing functional antibody responses.

## Results

### Monovalent XBB.1.5 mRNA booster elicited more robust and durable binding and neutralizing antibody responses compared to the bivalent mRNA booster

We profiled serum antibody responses by systems serology ^31-33^ from a cohort of healthcare workers (Table 1) vaccinated with the 2022-2023 bivalent mRNA vaccine (ancestral or wildtype (WT) + Omicron BA.5) or with the 2023-2024 monovalent mRNA vaccine (XBB.1.5). Serum samples were collected at baseline levels, ∼28 days post-immunization (peak immunogenicity), and ∼6 months post-immunization for both cohorts (**Figure 1A**, monovalent XBB.1.5 mRNA vaccination in purple, bivalent WT+BA.5 mRNA vaccination in green).

**Figure 1.**
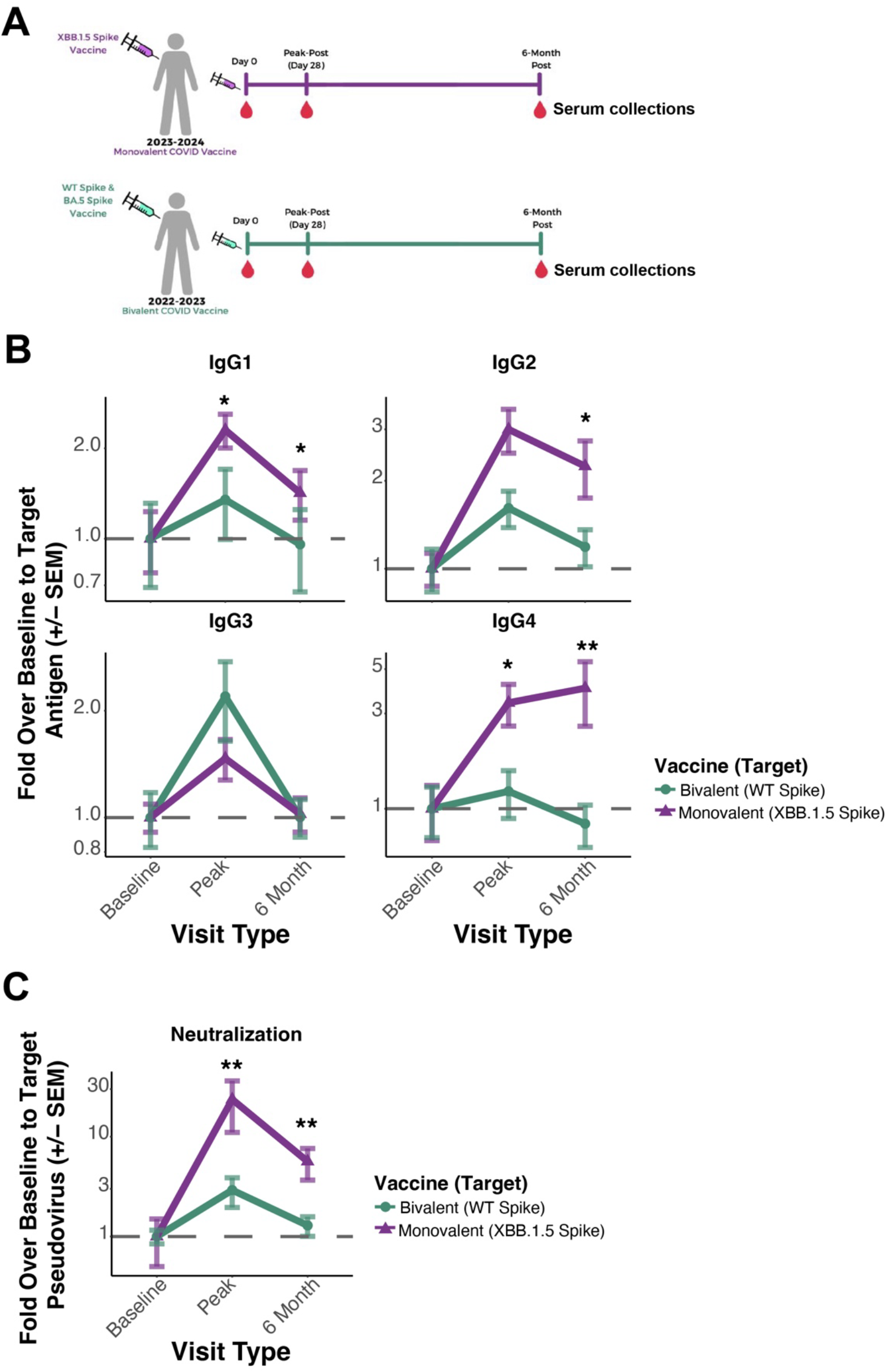
Comparison of antibody responses induced by the bivalent WT+BA.5 (2022-2023) or monovalent XBB.1.5 (2023-2024) mRNA COVID-19 boosters. (A) Scheme of cohort analysis. Participants were vaccinated with the monovalent XBB.1.5 (purple) or the bivalent WT+BA.5 (green) mRNA COVID-19 boosters. Serum samples were collected prior to vaccination (Day 0, or baseline), at peak immunogenicity post-vaccine (Day 28, or peak-post), and at 6 months post-immunization. Serum antibody responses were assessed by systems serology. (B) IgG subclass responses by vaccination cohort. Baseline (Day 0) antibody levels to the dominant target antigen (XBB.1.5 spike in purple, WT spike in green), peak (Day 28), and 6-month responses were quantified. Baseline values were standardized to 1 and shown on the y-axis are fold inductions over baseline (dotted line). Vaccine groupings legend is shown on the right. (C) Pseudovirus neutralizations to the target antigen (XBB.1.5 spike in purple, WT spike in green) were quantified similar to B. Statistical comparisons inter-group at the indicated timepoint were done using a Wilcoxon rank-sum test with a Bonferroni adjustment for multiple comparisons. The resulting q-values (Bonferroni adjusted p-value) were labeled as: * = q < 0.05, ** = q < 0.01, *** = q < 0.001.

We focused on IgG responses to the WT spike in individuals who received the bivalent mRNA boost (green) or to the XBB.1.5 spike in individuals who received the monovalent XBB.1.5 mRNA boost (purple). Spike-specific IgG1, the predominant subclass of IgG, was boosted significantly (2.6X-fold) higher for the monovalent vaccine compared to 1.3X-fold higher for the bivalent vaccine at peak response, and 1.4X fold higher at 6 months compared to a return to baseline for the bivalent recipients (**Figure 1B**, upper left; q-values = 0.02 and 0.015 for peak and 6 months post boost, respectively; two-sided Wilcoxon rank-sum test with Bonferroni adjustments for multiple comparisons). Similar differences were observed for spike-specific IgG2 and IgG4 responses, which were induced more robustly by the monovalent XBB.1.5 mRNA vaccine, particularly at month 6. In contrast, spike-specific IgG3 was induced similarly by both vaccines, potentially due to low absolute levels of IgG3 responses (**Figure 1B, Supplementary Figure 1A, B**). Differences in IgG1 responses to BA.5 following the bivalent mRNA boost compared with IgG1 responses to XBB.1.5 following the monovalent XBB.1.5 mRNA boost were even more pronounced (**Supplementary Figures 1C, 2A**).

We next looked at neutralizing antibody (NAb) responses generated by the XBB.1.5 monovalent and the WT/BA.5 bivalent mRNA vaccines to their target antigens. After vaccination, NAb titers increased 23X-fold to XBB.1.5 with the monovalent XBB.1.5 mRNA booster compared to a 2.9X-fold increase in NAb titers to the WT virus with the bivalent mRNA booster at peak immunogenicity, and 5.6X fold at 6 months compared to 1.3X fold for the bivalent recipients (**Figure 1C;** q-values = 0.0011 and 0.0026 for peak and 6 months post boost, respectively, two-sided Wilcoxon rank-sum test with Bonferroni adjustments for multiple comparisons). Similar results were observed for NAb titers to BA.5 with the bivalent mRNA booster (**Supplementary Figure 2B**).

These results suggest a more robust and durable antibody response to the target antigen by the monovalent XBB.1.5 mRNA booster compared to either target antigen of the bivalent mRNA booster. These results were not impacted by higher levels of infection-acquired immunity in the monovalent recipients (**Supplementary Figure 3**).

### Antibody correlations induced by the monovalent XBB.1.5 booster are broad and sustained

We next assessed how antibody correlations and humoral immune response architecture was shaped by the monovalent XBB.1.5 mRNA booster compared to the bivalent mRNA booster. We plotted high-confidence correlation cord diagrams for binding antibodies, Fc gamma receptor (FcγR) binding antibodies, Fc alpha receptor (FcαR) binding antibodies, and antibody Fc functions, including NAb responses, antibody-dependent cellular phagocytosis (ADCP), and antibody-dependent complement deposition (ADCD). In this analysis, only positive Spearman correlations were plotted (R > 0.5) that were significant after false discovery rate adjustments (p < 0.05 post correction).

In recipients of the bivalent mRNA booster, high-confidence correlations among various antibody parameters were limited at peak immunity (**Figure 2A**, top) and were further reduced after 6 months (**Figure 2A**, bottom). This was in contrast to multiple correlations among antibody parameters in recipients of the monovalent XBB.1.5 mRNA booster. In particular, binding and FcγR and FcαR-binding antibodies were directly correlated with neutralization and Fc effector functions at peak immunity (**Figure 2B**, top). After 6 months, much of the humoral immune response architecture was maintained, including numerous correlations between neutralization and binding and FcαR and FcγR-binding antibodies (**Figure 2B**, bottom). These results indicate that antibody responses to the 2022-2023 bivalent mRNA booster were largely disconnected, whereas antibody responses to the 2023-2024 monovalent XBB.1.5 mRNA booster were highly interconnected with many more correlations among various antibody features, suggesting more polyfunctional antibody responses.

**Figure 2.**
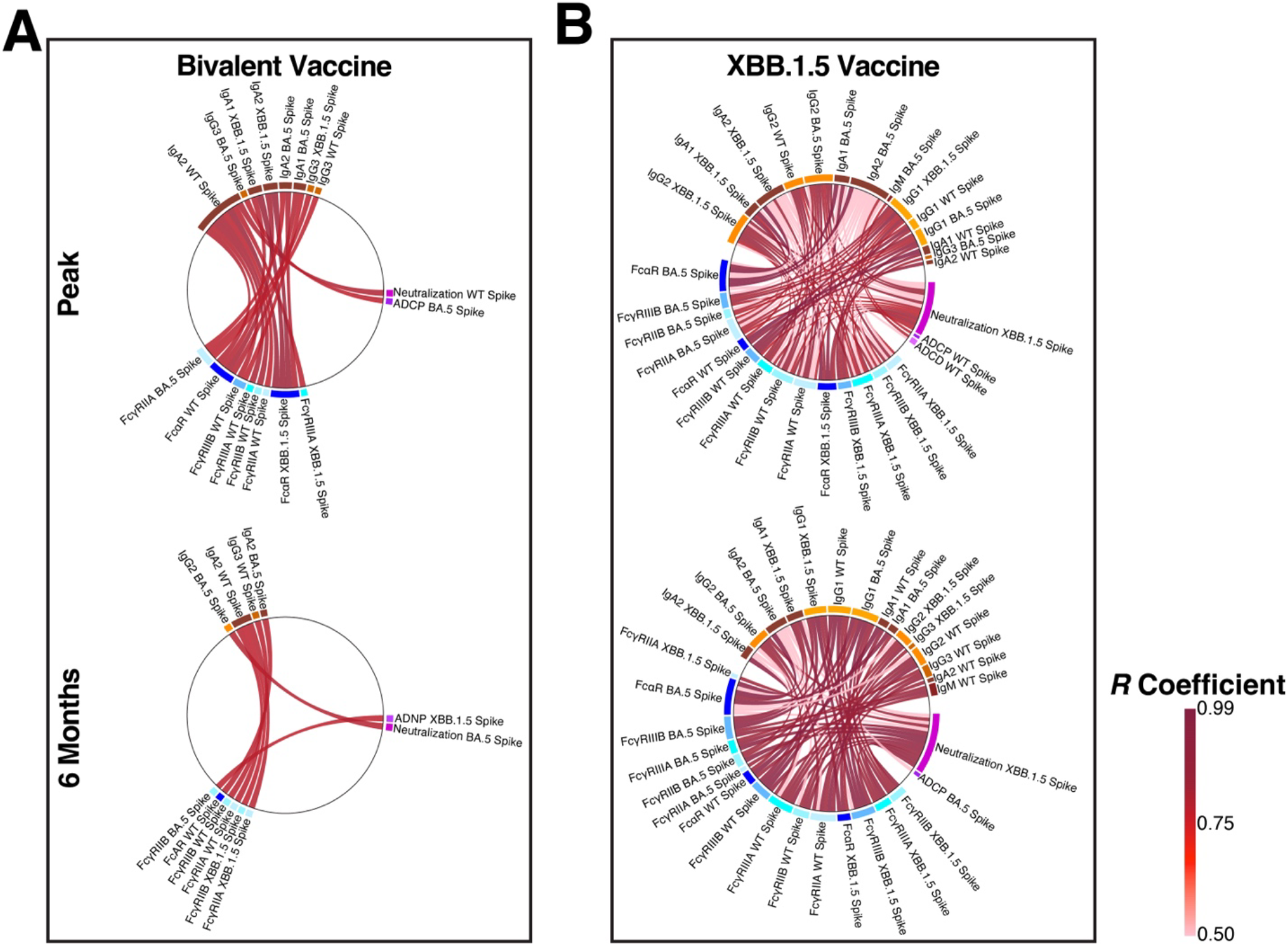
Antibody correlations and humoral immune response architecture induced by the bivalent WT+BA.5 or monovalent XBB.1.5 mRNA COVID-19 boosters. (A) Spearman’s correlation coefficients were calculated for pairwise comparisons for all systems serology assays in individuals who received the bivalent mRNA booster. Correlations whose false discovery rate (FDR) adjusted p-value < 0.05 were selected and plotted on a cord diagram for recipients of the 2022-2023 bivalent mRNA booster at peak (top) and 6-month post-vaccination responses. Binding antibody features are labeled next to their corresponding orange blocks, FcγR and FcαR-binding antibody features are labeled next to their corresponding blue blocks, and functional outputs are labeled next to their corresponding purple blocks. The size of the block is proportional to the number of directly co-correlated features. (B) Same as A, but in individuals who received the monovalent XBB.1.5 mRNA booster. Only pairings whose Spearman’s correlation coefficient was > 0.05 and FDR-adjusted p-value < 0.05 are shown. Legend for cord coloring is shown in the bottom right.

## Discussion

The spike protein of SARS-CoV-2 is the target of antibody-based recognition. As such, it is under selective pressure to evade previously established immunity while still retaining its ability to bind to ACE2 and promote infection. This has necessitated the deployment of updated boosters each year to address new circulating variants. In this study, we profiled antibody responses and correlations at peak immunogenicity and at 6 months post-immunization with the 2022-2023 bivalent mRNA booster and the 2023-2024 monovalent XBB.1.5 mRNA booster. The monovalent XBB.1.5 mRNA booster elicited a more robust and durable antibody response than the bivalent mRNA booster with more correlations among various antibody parameters, suggesting a more interconnected and functional antibody response.

It has previously been shown that NAb titers correlate with protection against SARS-CoV-2 ^10^. We recently reported that serum NAb responses in non-human primates were a strong predictor of protection against SARS-CoV-2 Omicron, while Fc functional antibody responses were a strong predictor of protection in the lower respiratory tract ^34^. FcγR- and FcαR-binding antibodies are frequently associated with effector function, but they also potentiate neutralization ^35,36^. The observation that both neutralizing and non-neutralizing antibody responses were highly coordinated following the monovalent XBB.1.5 mRNA booster suggests that the vaccine effectively induced antibodies with both Fab and Fc functional properties.

There are several potential explanations for why the monovalent XBB.1.5 mRNA boosters outperformed the bivalent mRNA boosters. First, the bivalent mRNA vaccine encoded for the WT spike in addition to BA.5, and the presence of the WT spike likely recalled WT-specific B cell responses by immune imprinting, which may have suppressed BA.5-specific responses ^27^. Second, population immunity in fall 2023 was higher than in fall 2022, which may have facilitated responses to the monovalent XBB.1.5 vaccine. We suspect that both the monovalent composition of the XBB.1.5 mRNA booster as well as the higher levels of baseline population immunity likely contributed to the more robust and durable antibody responses with this vaccine.

In summary, the monovalent XBB.1.5 mRNA booster yielded a robust, durable, and functionally interconnected antibody response for at least 6 months post-vaccination. These data support the continued deployment of monovalent rather than bivalent mRNA boosters that include and recall ancestral antigens.

## Methods

### Experimental model and participant details

The primary immunologic endpoint for recipients of the 2022-2023 bivalent vaccine were ancestral/wild type (WT) and Omicron BA.5 spike. The primary immunologic endpoint for recipients of the 2023-2024 monovalent booster was XBB.1.5 spike.

All participants were enrolled as a part of the Massachusetts Consortium on Pathogen Readiness (MassCPR) with informed consent. A demographic table of the two cohorts is shown in **Table S1**. Serum samples were obtained prior to receipt of the COVID-19 booster, at 28 days post-booster, and at 6 months post-booster. Booster vaccines were mRNA-based. For the bivalent vaccine, participants received the booster between September – December 2022. For the monovalent vaccine, participants received the booster between August – October 2023.

### Antigens

All antigens in this study are listed in the **Key Resources Table**. Antigens were obtained as lyophilized powder and were reconstituted in water to a final concentration of 0.5 mg/mL. Carbodiimide-NHS ester-coupling chemistry was employed to couple each antigen to its own microsphere region as previously described ^37^.

### Immunoglobulin isotype and Fc receptor binding

Levels of antigen-specific antibody isotypes, subclasses, and Fc receptor binding were determined with Luminex MagPlex microspheres (Luminex Corp, Austin, TX, USA) in technical replicates as previously described ^31-33^. Dilution curves identified each antigen’s linear range of detection in dilutions of human serum samples via secondary staining. Based on these results, antigen-coupled beads were incubated with diluted heat-inactivated serum (1:100 for IgG4, IgA1, IgA2, and IgM, 1:300 for Total IgG, IgG1, IgG2, IgG3, and FcαR, and 1:800 for FcγRIIA, FcγRIIB, FcγRIIIA, and FcγRIIIB) in 384-well plates (Greiner Bio-One, Frickenhausen, Germany) at 4°C overnight. The plates were washed to remove unbound antibodies and incubated for 1 hour with PE-conjugated secondary antibodies (1:100 for IgM, IgA1, and IgA2, 1:200 for Total IgG, IgG1, IgG2, IgG3, and IgG4). Fc receptors were biotinylated and bound to Streptavidin-R-Phycoerythrin (Agilent, Santa Clara, CA, USA) at a 1:1000 dilution. Remnant secondary reagent was removed by washing and the relative concentration of antibody per antigen was measured on the Luminex xMAP INTELLIFLEX DR-SE (Thermo Fisher, Waltham, MA, US).

### Evaluation of antibody-mediated functions

A flow cytometry-based monocyte phagocytic assay, antibody-dependent cellular phagocytosis (ADCP) was performed with custom synthesized fluorescent NeutrAvidin beads (Thermo Fisher, Waltham, MA, US) as previously described. Target antigens were biotinylated and conjugated to the beads. SARS-CoV-2 WT Spike and BA.5 Spike were conjugated to yellow-green microspheres, BQ.1.1 Spike and XBB.1.5 Spike were conjugated to scarlet microspheres, and Ebola virus glycoprotein (GP) was conjugated to red microspheres. Bead-conjugated WT Spike, BQ.1.1 Spike, and Ebola GP were combined in one set and BA.5 Spike, XBB.1.5 Spike, and Ebola GP were combined in another. These sets, and their technical repeats, were plated in 96-well U-bottom plates (Greiner Bio-One, Frickenhausen, Germany) with diluted serum (1:100) and incubated at 37°C and 5% CO_2_ for 2 hours. The plates were washed with 1X PBS and 25,000 cells/well of THP-1 monocytes were incubated overnight under the aforementioned conditions. Assay cells were fixed with 4% paraformaldehyde (PFA) before microsphere uptake readouts (“phagoscore”) were quantified using the iQue Screener Plus (Sartorius, Göttingen, Germany).

Antibody-dependent complement deposition (ADCD) was done as previously described ^37,38^. Briefly, heat-inactivated serum was diluted 1:10 and incubated with antigen-coated magnetic microspheres at 37°C for 2 hours with continuous shaking to bind antibodies to antigens. Beads were washed 3X to remove unbound material and guinea pig complement (Cedarlane, Burlington NC, USA) was added to the beads for 20 minutes at 37°C with continuous shaking. Complement deposition was stopped and quantification of complement deposition was done using anti-guinea pig C3-FITC (MP Biomedicals, Irvine CA, USA). Fluorescence for each microsphere was quantified on the iQue Flow Cytometer (Sartorius, Cambridge MA, USA). All samples were done in technical duplicates and means for each sample were reported.

Luciferase-based neutralization assays were performed as previously described against pseudoviruses containing the designated spike gene ^20^.

### Statistical Analysis

All assays were done in technical replicates for each sample. The mean was then taken for each sample for each assay. Baseline mean values were quantified and fold inductions relative to each group’s baseline mean were quantified. For statistical comparisons, an initial Wilcox test was performed followed by a Bonferroni correction for the number of comparisons made. All plots and statistics were done using ggplot 2 in R Studio V. 2023.03.0+386. For all comparisons, * indicates a q-value (Bonferroni-corrected p-value). Non-significant comparisons were left unlabeled.

For correlation cord diagrams, Spearman’s correlation coefficients (R) were quantified along with false discovery rate (FDR) corrected p-values for each pairing. Only positive Spearman’s correlation coefficients > 0.5 and FDR-corrected p-value < 0.05 were plotted. Cord diagrams were generated using R studio V 2023.03.0+386.

## Acknowledgments

This work was supported by the Massachusetts Consortium for Pathogen Readiness (MassCPR), the U01CA260476-02, and the 1P01AI168347–01 to R.P.M. We also thank Nancy Zimmerman, Mark and Lisa Schwartz, an anonymous donor (general financial contribution to the Ragon Institute), and Terry and Susan Ragon for their donations to the Ragon Institute.

## Author Contributions

Conceptualization: S.E.B., K.S.L., and R.P.M.

Data curation: S.E.B., K.S.L., and R.P.M.

Formal analysis: S.E.B., K.S.L., Q.W., X.T., and R.P.M.

Funding acquisition: R.P.M.

Investigation: S.E.B., K.S.L., R.B., Q.W., X.T., and R.P.M.

Methodology: S.E.B., K.S.L., R.B., Q.W., X.T., and R.P.M.

Project administration: Q.W., X.T., and R.P.M.

Resources: R.P.M.

Software: R.P.M

Supervision: Q.W., X.T. and R.P.M.

Validation: S.E.B., K.S.L., R.P.M.

Visualization: S. E. B. K.S.L., and R. P. M.

Writing–original draft: S. E. B., K.S.L., and R.P.M.

## Key Resources Table

| REAGENT or RESOURCE | SOURCE | IDENTIFIER |
| --- | --- | --- |
| **Antibodies** |  |  |
| Mouse Anti-Human IgG Fc-PE (JDC-10) | SouthernBiotech | 9040-09 |
| Mouse Anti-Human IgG1 Hinge-PE (4E3) | SouthernBiotech | 9052-09 |
| Mouse Anti-Human IgG2 Fc-PE (31-7-4) | SouthernBiotech | 9060-09 |
| Mouse Anti-Human IgG3 Hinge-PE (HP6050) | SouthernBiotech | 9210-09 |
| Mouse Anti-Human IgG4 Fc-PE (HP6025) | SouthernBiotech | 9200-09 |
| Mouse Anti-Human IgA1-PE (B3506B4) | SouthernBiotech | 9130-09 |
| Mouse Anti-Human IgA2-PE (A9604D2) | SouthernBiotech | 9140-09 |
| Mouse Anti-Human IgM-PE (SA-DA4) | SouthernBiotech | 9020-09 |
| Goat Anti-Guinea Pig Complement C3 FITC | MP Biomedicals | 0855385 |
| **Chemicals, Peptides, and Recombinant Proteins** |  |  |
| SARS-CoV-2 Alpha RBD | Sino Biological | 40592-V08H82 |
| SARS-CoV-2 Alpha Spike | Sino Biological | 40589-V08B6 |
| SARS-CoV-2 BA.1 RBD | Sino Biological | 40592-V08H121 |
| SARS-CoV-2 BA.1 Spike | Sino Biological | 40589-V08H26 |
| SARS-CoV-2 BA.2.86 Spike | Sino Biological | 40589-V08H58 |
| SARS-CoV-2 BA.5 RBD | Sino Biological | 40592-V08H130 |
| SARS-CoV-2 BA.5 Spike | Sino Biological | 40589-V08H32 |
| SARS-CoV-2 Beta RBD | Sino Biological | 40592-V08H59 |
| SARS-CoV-2 Beta Spike | Sino Biological | 40589-V08B7 |
| SARS-CoV-2 BQ.1.1 RBD | Sino Biological | 40592-V08H143 |
| SARS-CoV-2 BQ.1.1 Spike | Sino Biological | 40589-V08H41 |
| SARS-CoV-2 D614G Spike | Sino Biological | 40589-V08H8 |
| SARS-CoV-2 Delta RBD | Sino Biological | 40592-V08H90 |
| SARS-CoV-2 Delta Spike | Sino Biological | 40589-V08H10 |
| Ebola Virus Glycoprotein | Sino Biological | 40459-V08H |
| SARS-CoV-2 EG5.1 RBD | Sino Biological | 40592-V08H151 |
| SARS-CoV-2 EG5.1 Spike | Sino Biological | 40589-V08H55 |
| SARS-CoV-2 Gamma RBD | Sino Biological | 40592-V08H86 |
| SARS-CoV-2 Gamma Spike | Sino Biological | 40589-V08H23 |
| HCMV Glycoprotein B | Sino Biological | 10202-V08H1 |
| Human Coronavirus HKU1 Spike | Sino Biological | 40606-V08B |
| SARS-CoV-2 JN.1 RBD | Sino Biological | 40592-V08H155 |
| SARS-CoV-2 JN.1 Spike | Sino Biological | 40589-V08H59 |
| MERS-CoV Spike | Sino Biological | 40069-V08B |
| SARS-CoV-2 WT Nucleocapsid | Sino Biological | 40588-V08B |
| HCoV-NL63 Spike | Sino Biological | 40604-V08B |
| Human Coronavirus HCoV-OC43 Spike | Sino Biological | 40607-V08B |
| SARS-CoV-1 Spike | Sino Biological | 40634-V08B |
| SARS-CoV-2 WT NTD | Sino Biological | 40591-V49H |
| SARS-CoV-2 WT RBD | Sino Biological | 40592-V08H |
| SARS-CoV-2 WT S1 | Sino Biological | 40591-V08H |
| SARS-CoV-2 WT S2 | Sino Biological | 40590-V08B |
| SARS-CoV-2 WT Spike | Sino Biological | 40589-V08H4 |
| SARS-CoV-2 XBB.1.16 RBD | Sino Biological | 40592-V08H136 |
| SARS-CoV-2 XBB.1.16 Spike | Sino Biological | 40589-V08H48 |
| SARS-CoV-2 XBB.1.5 RBD | Sino Biological | 40592-V08H146 |
| SARS-CoV-2 XBB.1.5 Spike | Sino Biological | 40589-V08H45 |
| Human soluble FcγRIIA | Duke University | Custom Order |
| Human soluble FcγRIIB | Duke University | Custom Order |
| Human soluble FcγRIIIA | Duke University | Custom Order |
| Human soluble FcγRIIIB | Duke University | Custom Order |
| Human soluble FcαR | Duke University | Custom Order |
| EZ-Link Sulfo NHS-LC-LC Biotin | Thermo Fisher | A35358 |
| Sulfo-NHS (N-hydroxysulfosuccinimide) | Thermo Fisher | A39269 |
| Pierce EDC | Thermo Fisher | A35391 |
| BirA500: BirA biotin-protein ligase standard reaction kit | Avidity LLC | BirA500 |
| Streptavidin-R-Phycoerythrin | Agilent | PJ31S |
| **Experimental Models: Cell Lines** |  |  |
| THP-1 monocytes | ATCC | RRID: CVCL_0006 |
| **Software and Algorithms** |  |  |
| iQue Forecyt 9.1 | Sartorius | 60028 |
| R Studio V 4.0 | R Project for Statistical Computing | RRID:SCR_000432 |
| **Other** |  |  |
| iQue Screener Plus | Sartorius | 11811 |
| xMAP INTELLIFLEX DR-SE System | Thermo Fisher | APX2021 |
| MagPlex Microspheres | Diasorin | MC12001-01<br>(Cataloged by region) |
| Yellow-Green Fluorescent NeutrAvidin Microspheres | Thermo Fisher | Custom Synthesis |
| Red Fluorescent NeutrAvidin Microspheres | Thermo Fisher | Custom Synthesis |
| Scarlet Fluorescent NeutrAvidin Microspheres | Thermo Fisher | Custom Synthesis |
| Low-Tox Guinea Pig Complement, Lyophilized | Cedarlane | CL4051 |
| 384-well HydroSpeed Plate Washer | Tecan | 30190112 |
| Zebra-Spin Desalting and Chromatography Columns | Thermo Fisher | 89882 |

