## Supplementary Information for "Monovalent XBB.1.5 mRNA Vaccine Recalls a More Durable and Coordinated Antibody Response to SARS-CoV-2 Spike than the Bivalent WT/BA.5 mRNA Vaccine"

Supplementary information includes Supplementary Table 1 and Supplementary Figures 1-3 and their corresponding Figure Captions.

### Table S1

| | Cohort | |
|---|---|---|
| Vaccine | Bivalent (WT + BA.5) | Monovalent (XBB.1.5) |
| N | 42 | 31 |
| Mean age and range | 42 (24-70) | 37 (22-73) |
| Sex | 70% Female | 81% Female |
| | 30% Male | 19% Male |
| Manufacturer | Pfizer/BioNTech (15) | Pfizer/BioNTech (13) |
| | Moderna (27) | Moderna (18) |

**Table S1. Cohort background information**

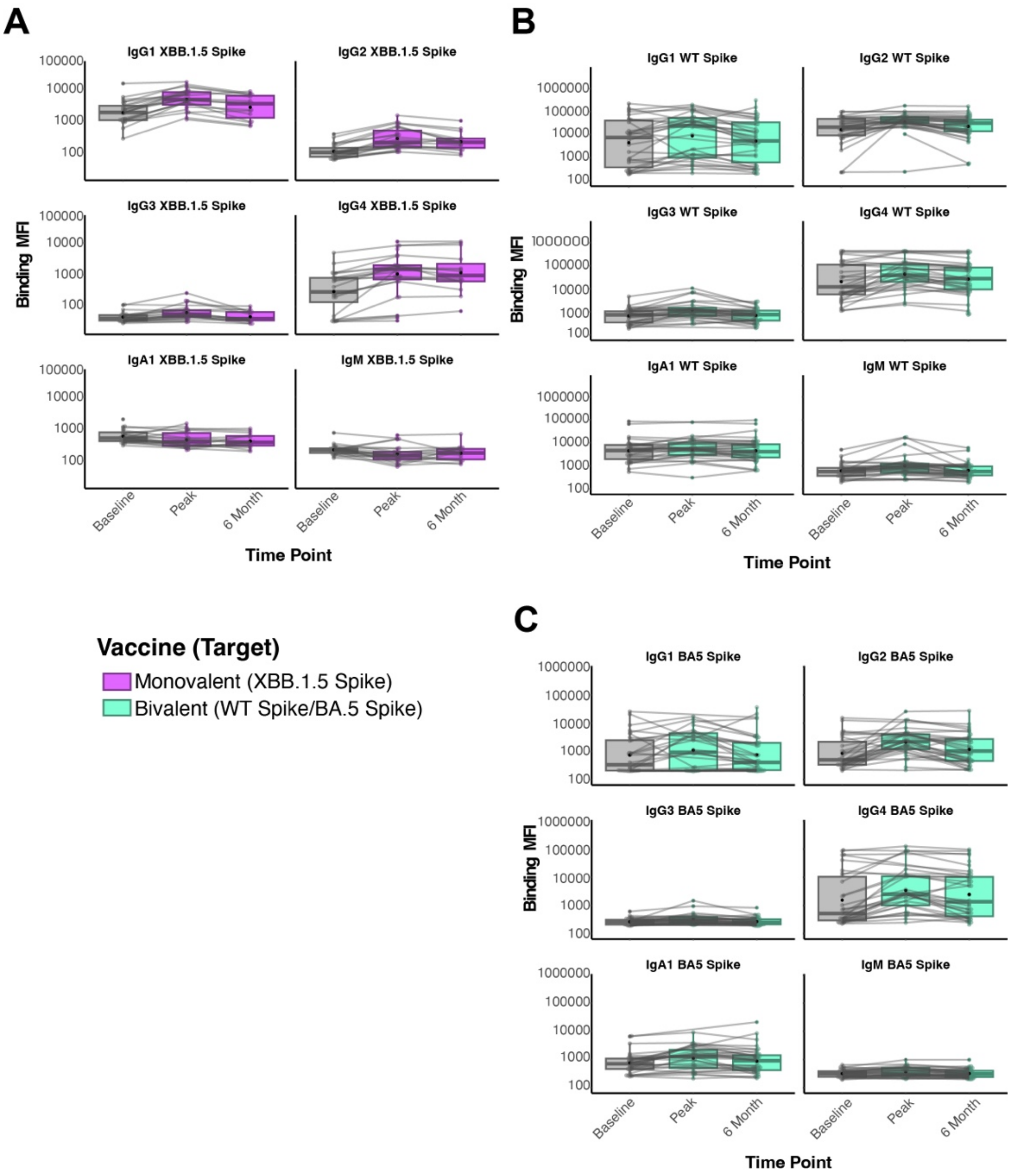


**Supplementary Figure 1. Unadjusted values for vaccine responses.** (A) Raw binding mean fluorescence intensity (MFI) for the indicated antibody subclasses and isotypes to XBB.1.5 spike in individuals who received the monovalent XBB.1.5 mRNA booster. (B) Same as A, but for responses to WT spike in individuals who received the bivalent mRNA booster. (C) Same as A, but for responses to BA.5 spike in individuals who received the bivalent mRNA booster. The color legend is shown on the bottom left. Shown are individual data points representing individual participants with lines connecting their responses over time. Box and whisker plots are superimposed showing the median, interquartile ranges, and 95% confidence intervals.

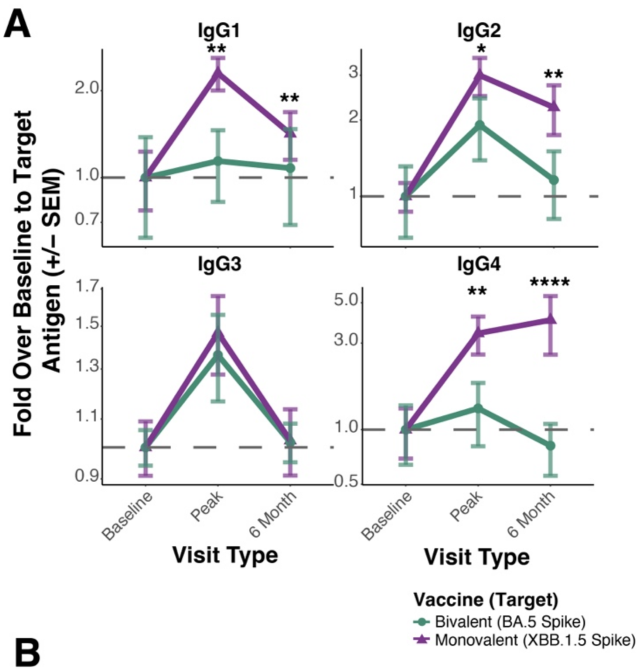


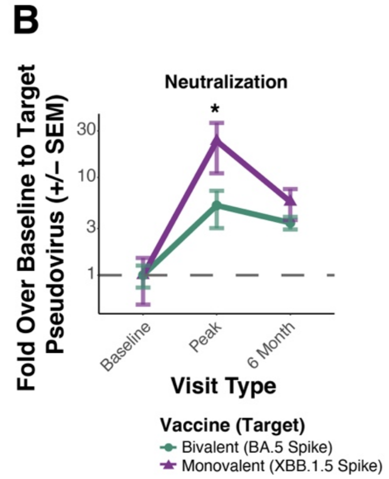


**Supplementary Figure 2. Comparison of antibody responses induced by the bivalent WT+BA.5 (2022-2023) or monovalent XBB.1.5 (2023-2024) mRNA COVID-19 boosters.** (A) IgG subclass responses by vaccination cohort. Baseline (Day 0) antibody levels to the target antigen (XBB.1.5 spike in purple, BA.5 spike in green), peak (Day 28), and 6 month responses were quantified. Baseline values were standardized to 1 and shown on the y-axis are fold inductions over the baseline (dotted line). Vaccine groupings legend is shown on the right. (C) Pseudovirus neutralizations to the target antigen (XBB.1.5 spike in purple, BA.5 spike in green) were quantified similarly to B. Statistical comparisons inter-group at the indicated timepoint were done using a Wilcoxon rank-sum test with a Bonferroni adjustment for multiple comparisons. The resulting q-values (Bonferroni adjusted p-value) were labeled as: * = q < 0.05, ** = q < 0.01, *** = q < 0.001.

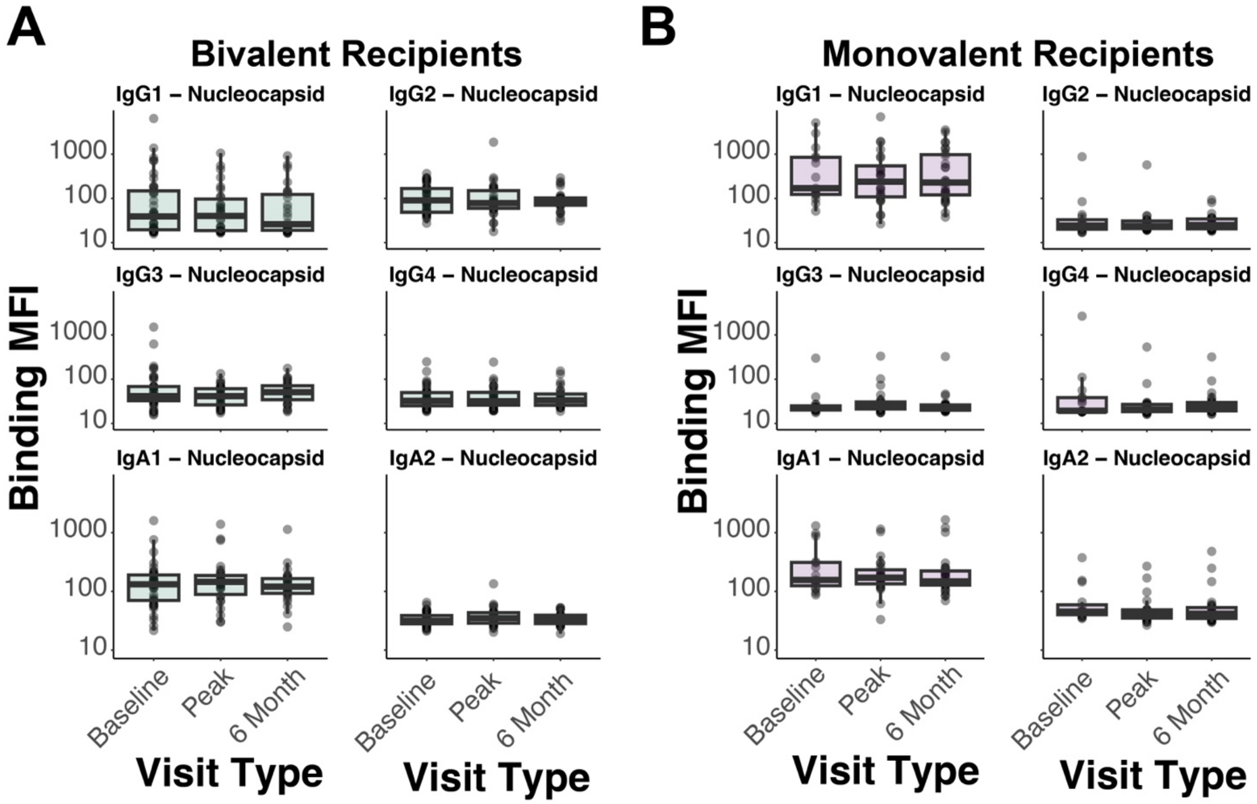


**Supplementary Figure 3. Infection-acquired immunity did not shape immune profiles for the two cohorts.** (A) Raw binding MFI values to nucleocapsid (an indicator of infection) are shown for antibody subclasses and isotypes in individuals who received the bivalent mRNA booster. (B) Same as A, but in individuals who received the monovalent XBB.1.5 mRNA booster. Box and whisker plots are superimposed showing the median, interquartile ranges, and 95% confidence intervals. No significant differences relative to baseline were identified after Wilcoxon rank-sum test with a Bonferroni adjustment for multiple comparisons.
